# Research advance: Unexpected plasticity in the life cycle of *Trypanosoma brucei*

**DOI:** 10.1101/2025.08.20.671235

**Authors:** Carina Praisler, Jaime N. Lisack, Anna Sophie Kreis, Laura Hauf, Johanna Krenzer, Fabian Imdahl, Markus Engstler

**Affiliations:** Department of Cell and Developmental Biology, Biocenter, Julius-Maximilians-Universitaet, Wuerzburg, Germany; Helmholtz Institute for RNA-based Infection Research, Helmholtz-Center for Infection Research, Wuerzburg, Germany

## Abstract

We have previously shown that the slender form of *Trypanosoma (T.) brucei* is able to infect teneral tsetse flies, develop to the first fly form, which is the procyclic form, and complete the life cycle in the insect vector (Schuster et al., 2021). Further, analysis of the transmission index (TI; defined as the number of salivary gland infections relative to the number of midgut infections) revealed a higher TI for slender as compared to stumpy forms under laboratory conditions, which included the addition of *N*-acetyl-glucosamine (NAG) to the infective bloodmeal.

These findings challenge the prevailing view of the life cycle, according to which only stumpy forms are considered infective to tsetse flies.

Here, we show that slender trypanosomes can infect both male and female tsetse flies, irrespective of their teneral status, in the absence of supplements in the bloodmeal.

Additionally, an RNA-sequencing time course was performed on both slender and stumpy cells during their transition into procyclic forms. This analysis revealed that slender and stumpy form trypanosomes remain transcriptionally distinct throughout differentiation into the procyclic form. Furthermore, while the protein associated with differentiation 1 (PAD1) remains essential for the transition, slender cells do not require expression of other hallmark stumpy form traits, such as cell cycle arrest or the shortening of their flagella or microtubule corset. Instead, slender trypanosomes are able to transition directly into procyclic forms.

Taken together, these findings demonstrate that while slender cells of *T. brucei* follow distinct routes to become the procyclic form, they are capable of infecting both teneral and non-teneral tsetse flies, thereby contributing to the transmission and spread of these African parasites.

## Introduction

In the bloodstream of a mammalian host, two main forms of *Trypanosoma (T.) brucei* can be observed, the long slender and the short stumpy bloodstream form (bsf).

The slender cell is proliferative whereas the stumpy cell has undergone cell-cycle arrest in G1/G0 phase. During division, slender cells release peptidases, most significantly oligopeptidase B and metallocarboxypeptidase 1(Tettey et al., 2022), generating a pool of oligopeptides in the blood and tissues around them. These essential peptides, together with possible other unknown components, are part of the molecular cocktail collectively called the Stumpy Induction Factor (SIF) (Vassella et al., 1997; Reuner et al., 1997; Bossard et al., 2013; Moss et al., 2015; Rojas et al., 2019). After exceeding a certain threshold, SIF causes the proliferative slender cells to change into the cell-cycle arrested stumpy forms (Vassella et al., 1997; Reuner et al., 1997). This transition is accompanied by shortening of the flagellum, cell-cycle arrest and other molecular changes, such as the remodelling of the mitochondrion to a cristate structure and expression of Krebs cycle enzymes. The trypanosomes also begin to express the Protein Associated with Differentiation 1 (PAD1), a member of the carboxylate-transporter protein family, on their surface. PAD1 functions as a transducer of the signal that triggers differentiation into the procyclic form, the first fly form that develops in the tsetse midgut (Reuner et al., 1997; Dean et al., 2009).

Cell-cycle arrest is lethal for stumpy forms, as they die, if they are not taken up by the tsetse fly within a few days. Thus, their purpose in the life cycle of *T. brucei* has long been debated, with the two most accepted theories being:

1. Stumpy forms are needed for the regulation of the parasitaemia in the host.
2. Due to pre-adaptation for life in the insect vector, they are the only form able to infect the tsetse fly (Robertson, 1912; Wijers & Willett, 1960; Fenn & Matthews, 2007; MacGregor et al., 2012; Matthews et al., 2015; Silvester et al., 2017).

In 2021, our laboratory reported unexpected plasticity in the life cycle of *T. brucei*. We found that even a single slender cell can infect the tsetse fly without exhibiting morphological and biochemical manifestations that define a stumpy cell (Schuster et al., 2021).

Instead, slender cells turn on the essential PAD1 pathway (Dean et al., 2009) and transit directly into procyclic forms - all whilst continuously dividing. We suggested that the ability of slender bloodstream forms to infect the tsetse fly vector would, at least in part, solve the transmission paradox, which refers to the low blood parasitaemia observed in chronically infected hosts and the small bloodmeal volume of tsetse flies, both of which make it unlikely that a stumpy form would be ingested (Capewell et al., 2019).

However, our original study was criticised (Matthews & Larcombe, 2022; Ngoune et al., 2025) for mainly four reasons:

(1) As routinely done in tsetse laboratories, we supplemented all infectious bloodmeals with the immune-suppressive chemical *N*-acetyl-glucosamine (NAG), which is known to enhance tsetse midgut infections for stumpy forms but has no effect on subsequent salivary gland infections (Peacock et al., 2012). It was argued that this treatment allowed slender forms to infect the fly.
(2) We used teneral flies (which have not yet taken a bloodmeal and younger than 3 days) for all infection experiments. This led to the question if the slender trypanosomes might just be able to infect young flies.
(3) The use of male flies is common practice for studying trypanosome infections. Nevertheless, could it be possible that slender trypanosomes can infect male but not female tsetse flies?
(4) We had shown that slender trypanosomes express the differentiation marker *pad1* while becoming procyclic. This was taken as a proof that slender cells must turn into stumpy cells before becoming procyclic.

To address the critique further than already done (Lisack et al., 2022), we have systematically conducted additional experiments. We found that slender form trypanosomes can indeed infect both teneral and non-teneral flies (i. e. flies that have taken at least one non-infectious bloodmeal and are more than 72 hours post eclosion (hpe)), even in the absence of immune suppressing chemicals.

Using an RNA-sequencing time course, we confirmed that slender forms are able to transition directly to procyclic forms without becoming stumpy forms.

Taken together these new findings further highlight the plasticity in the life cycle of *T. brucei* as well as the ability of slender trypanosomes to contribute to the spread of these parasites to new hosts.

## Results

It has been shown that *N*-acetyl-glucosamine (NAG) aids stumpy trypanosome infection of the fly midgut, while not influencing the number of subsequent salivary gland infections (Peacock et al., 2006, 2012).

To exclude the possibility that use of the lectin-inhibitor NAG rendered the tsetse fly midgut artificially permissive to slender trypanosome infections, teneral tsetse flies were infected with slender cells in either untreated blood or NAG-supplemented blood.

As described in Schuster *et al*., a stumpy marker cell line (NLS-GFP:PAD1 3’UTR) was used to confirm that less than 1% PAD1-positive cells were present in any slender culture.

Infection rates in the midgut and proventriculus showed negligible differences between non-supplemented and NAG-supplemented infections (Figure 1A and Supplementary Figure 1). Midgut infections reached 11.6% with and 9.2% without NAG (Figure 1A) while proventriculus infections occurred in 9.4% and 8.3%, respectively (Figure 1A). Salivary gland infections were observed in 3.6% of flies infected with and 0.9% without NAG, however, this difference was not statistically significant (p > 0.05) (Figure 1A).

**Figure 1:**
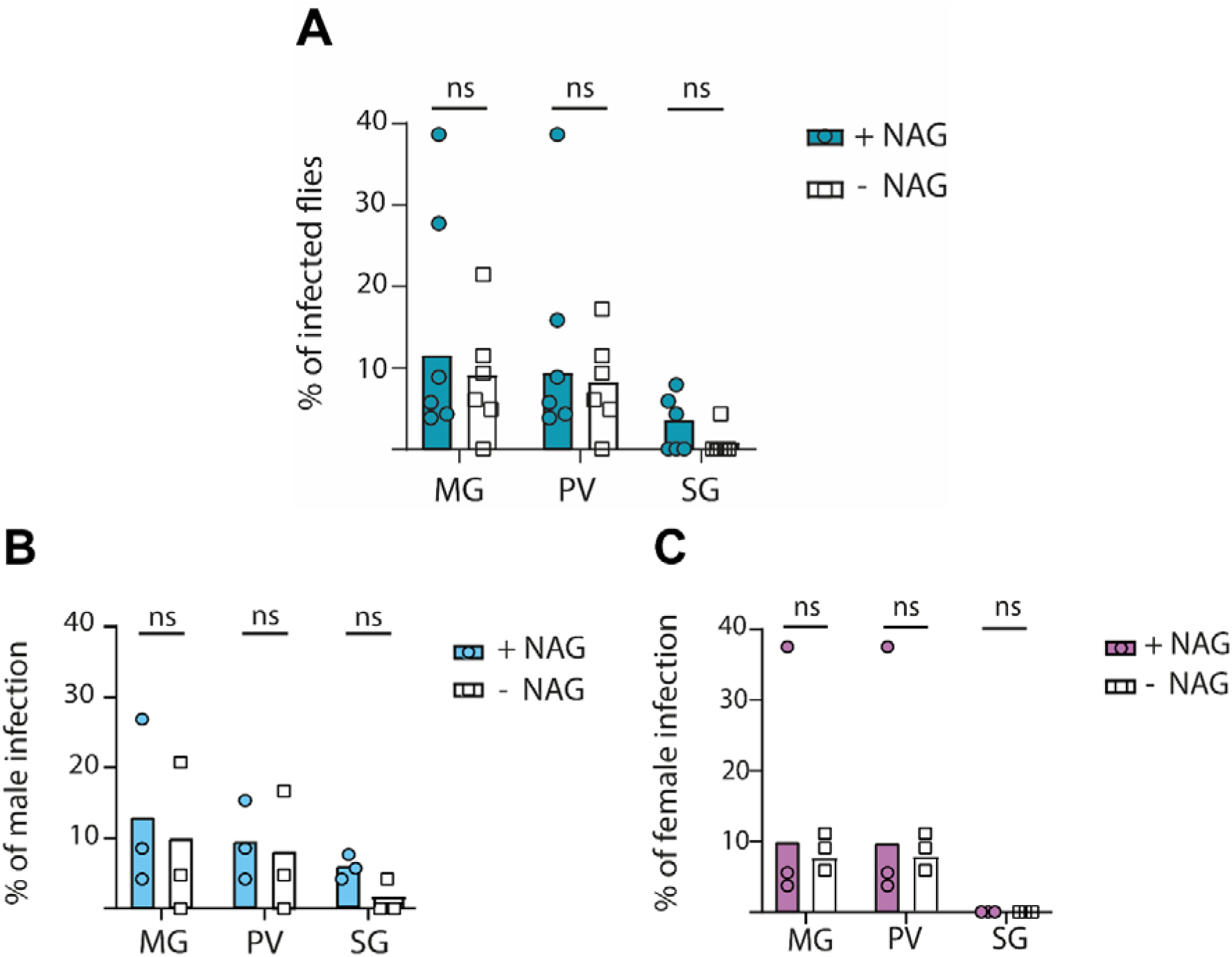
Infection rates (%) of slender *T. brucei* in teneral tsetse flies, with and without NAG supplementation. Flies of both genders were fed 4 slender cells per bloodmeal, with or without 60 mM *N*-acetyl-glucosamine (NAG). Slender cells were harvested and checked for PAD1 signal to confirm slender identity before infection (< 1% PAD1 positive). Infections were performed in triplicates with approximately 20 flies/replicate. The midgut (MG), proventriculus (PV) and salivary glands (SG) of all flies were dissected after 35 days to check for infection. Bar graphs show mean infection rates across replicates, with individual dots representing infection percentages per replicate. Fisher’s exact test was used on the mean infection rates to determine significance. ns = not significant (p > 0.05). Infection rates are shown for both sexes (A), male (B), and female flies (C).

The use of male flies is common practice for studying tsetse infections, as they show higher salivary gland infection rates than female flies (Jackson, 1949; Maudlin, 1990; Maudlin et al., 1991). However, because female flies have a longer lifespan, they cannot be disregarded as potential important contributors to parasite transmission in the wild (Maudlin et al., 1990). Therefore, the influence of fly sex on infection rates was assessed by using the same dataset as in Figure 1A (Figure 1B and C). Using NAG-supplemented blood, 8.0% of male flies exhibited midgut, 5.8% proventriculus and 3.6% salivary gland infections. In the absence of NAG, midgut, proventriculus, and salivary gland infections were observed in 5.5%, 4.6%, and 0.9% of male flies, respectively (Figure 1B). Fisher’s exact test revealed no significant differences in infection rates between NAG-supplemented and non-supplemented groups for male flies (Figure 1B).

Among female flies, 3.6% harboured midgut and 3.7% proventriculus infections, regardless of NAG supplementation (Figure 1C). No salivary gland infections were identified in female flies (Figure 1C). The total percentage of fly infections and their corresponding replicates are provided in Supplementary Figure 1.

In conclusion, NAG has a negligible effect on slender infections in tsetse flies and even low numbers of slender cells can infect flies without the aid of immune suppressing compounds.

Although infection rates for teneral flies are relatively low, commonly ranging between 10-30%, they are more susceptible to trypanosome infection than non-teneral flies (Peacock et al., 2012). Consequently, the use of teneral flies is standard practice in *T. brucei* infection experiments, and we have exclusively used them in our previous infections. It is possible, however, that in nature, older flies contribute more to the spread of trypanosomes than younger flies. As such, it was of interest to test if slender trypanosomes could also infect non-teneral flies (Van Hoof L. et al., 1937; Wijers, 1958; Otieno et al., 1983; Walshe et al., 2011; Matthews & Larcombe, 2022). Therefore, to ensure our flies would survive for at least 30 days to allow infection assessment, we defined our non-teneral flies as those infected between 144 and 168 hours post eclosion (hpe). These flies were fed two non-infectious bloodmeals prior to the third infectious bloodmeal, all spaced at least two days apart.

For these experiments, we again used the cell line containing the NLS-GFP:PAD1 3’UTR stumpy reporter and a cytoplasmic red fluorescent protein (td Tomato) (Reuter et al., 2023). To ensure pure populations (Schuster et al., 2021), fluorescence activated cell sorting (FACS) was performed to separate PAD1-positive (stumpy) from PAD1-negative (slender) cells. To confirm sorting success, populations were subsequently examined by fluorescence microscopy for PAD1 signal. Stumpy cells were also sorted to remove any slender cells and to keep conditions constant.

Additionally, the growth of sorted slender cells was monitored *in vitro* to ensure that sorting did not affect parasite fitness by causing any stress (Quintana et al., 2021) (Supplementary Figures 3 and 4). Only then did we proceed with infections, using either sorted slender or sorted stumpy cells to infect male and female, teneral or non-teneral flies. With the confidence of having pure slender or stumpy populations, we increased parasite concentrations to 1×10^6^ cells/ml for infection. Flies were dissected 30 days post infection and their organs examined for parasite presence.

Infection data show that slender trypanosomes were able to establish infections in non-teneral flies at rates comparable to stumpy forms (Figure 2A and Supplementary Figure 2). Midgut infections were detected in 6.4% of flies infected with slender and 6.6% of flies infected with stumpy cells. Similarly, 5.1% and 4.4% of flies exhibited proventriculus infections, and 3.8% and 1.5% salivary gland infections, respectively. Statistical analysis using Fisher’s exact test revealed no significant difference for any fly organ in slender or stumpy infections.

**Figure 2:**
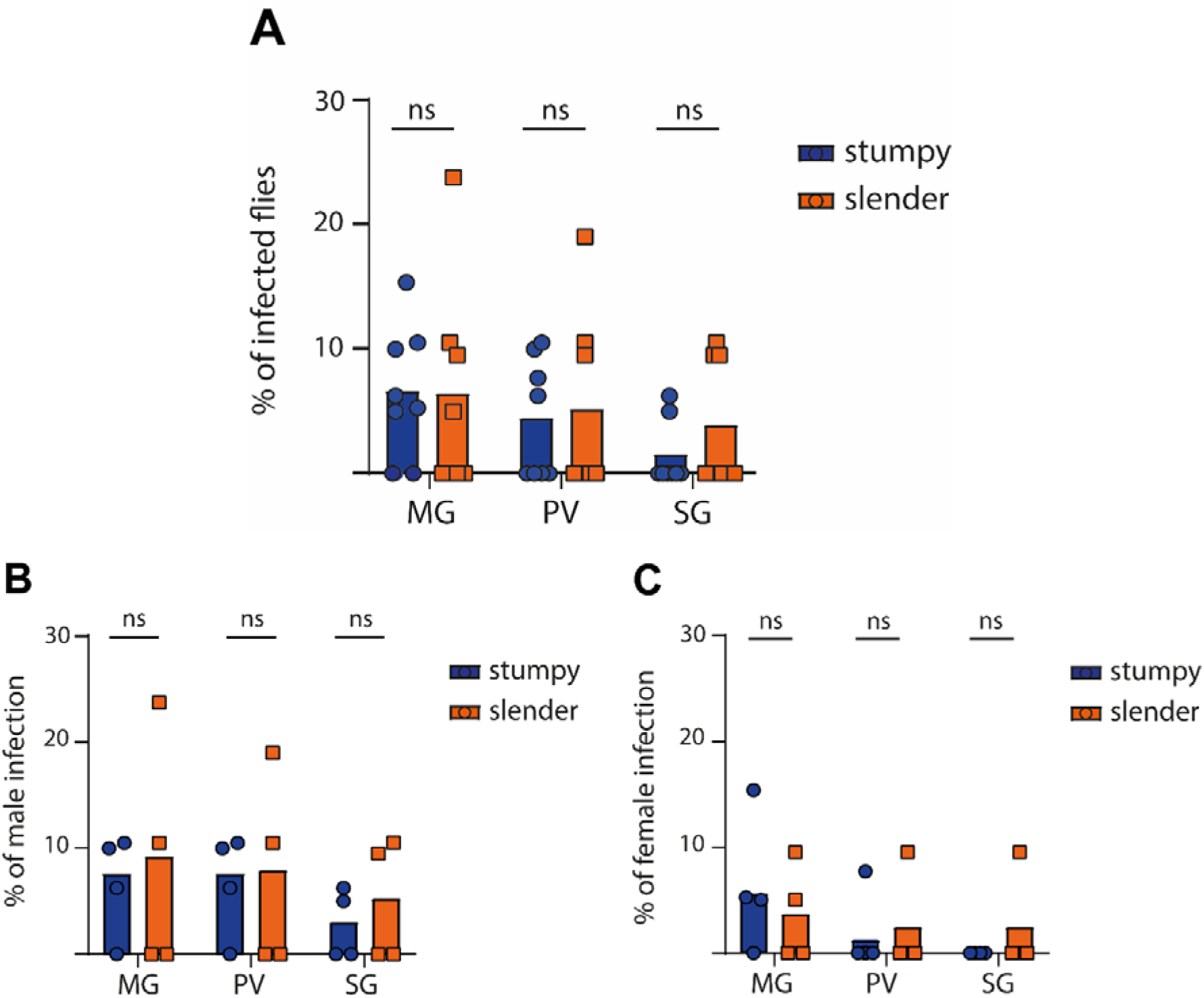
Infection rates (%) of slender and stumpy *T. brucei* in non-teneral tsetse flies. Non-teneral flies were infected between 144 and 168 hpe with either slender (orange) or stumpy (blue) parasites, after receiving two non-infectious bloodmeals beforehand. The tdTomato NLS-GFP:PAD1 3’UTR cell line enabled FACS to separate stumpy (PAD1 positive, GFP in nucleus) and slender (PAD1 negative, no nuclear fluorescence) prior to infection. Infections were performed in quadruplets with roughly 20 flies/replicate. Midgut (MG), proventriculus (PV), and salivary glands (SG) of all flies were dissected after 30-35 days to check for parasite presence. Bar graphs show mean infection rates across replicates, with individual dots representing infection percentages per replicate. Fisher’s exact test was used on the mean infection rates to determine significance; ns = not significant (p > 0.05). Infection rates are presented for both sexes (A), males (B), and females (C).

Consistent with published data on stumpy infections (Peacock et al., 2012), male flies showed the highest infection rates (Figure 2B). Importantly, we found that non-teneral female flies can also be infected with slender cells. Midgut infections were observed in 9.5% of non-teneral female flies infected with slender cells and 5.6% with stumpy cells (Figure 2C). Although overall numbers were low, it is noteworthy that only slender cells caused any salivary gland infections in non-teneral female flies (Figure 2C). A table of all percentages of fly infections, replicates, and sex can be found in Supplementary Figure 2.

We have previously shown that slender and stumpy forms both differentiate to procyclic forms with comparable kinetics (Schuster et al. 2021). To assess whether slender cells transcriptionally transition through a stumpy state or differentiate directly into the procyclic form, we performed RNA sequencing of slender and stumpy parasites throughout differentiation. For this, we used the same cell line as in Schuster *et al*. (2021), containing an EP1:YFP fusion protein and NLS-GFP:PAD1 3’UTR stumpy reporter. *In vitro* differentiation of both slender and stumpy cultures was induced by glucose depletion, addition of *cis*-aconitate, and temperature drop to 27°C (Mowatt & Clayton, 1987; Richardson et al., 1988; Roditi et al., 1989; Ziegelbauer et al., 1990; Matthews & Gull, 1994; Dean et al., 2009).

Slender or stumpy cells were collected in triplicates at five time-points throughout differentiation, 0, 8, 15, 24, and 72 hours (hrs), and subjected to paired-end sequencing (Figure 3). Hierarchical clustering identified two biological replicates as outliers, which were excluded from further analysis.

**Figure 3:**
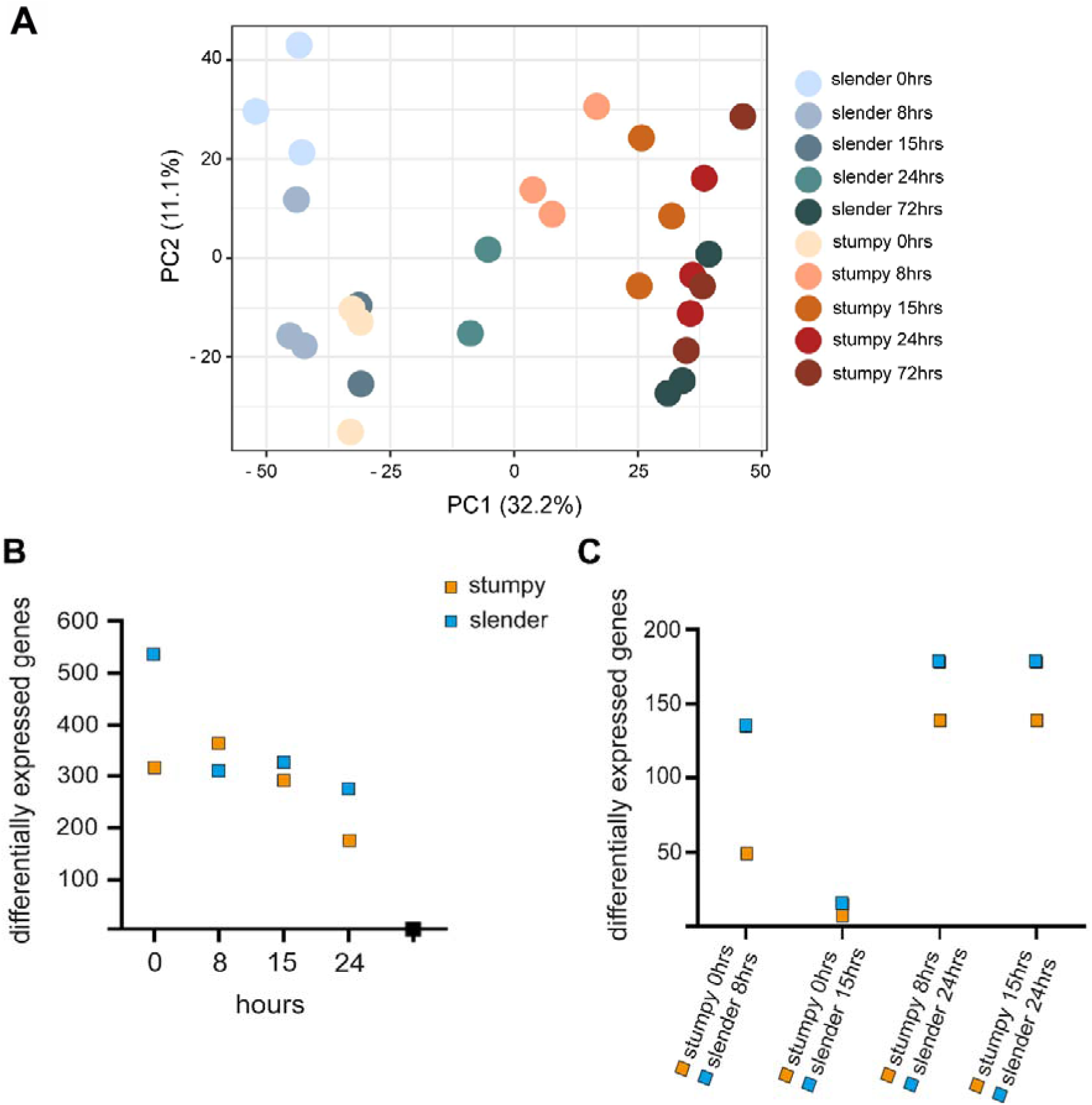
RNA sequencing of slender and stumpy trypanosomes differentiating into procyclic forms, performed in biological triplicates. A: Principal Component Analysis (PCA) showing the transcriptional progression to procyclic forms for slender (blue/green) and stumpy (orange/red) cells. The trajectories remain distinct until converging at 72hrs. B: Number of differentially expressed genes that are upregulated in slender compared to stumpy forms at corresponding time points during differentiation. Genes with an absolute log2FC > 2 and an adjusted p-value < 0.01 were classified as differentially expressed. Corresponding volcano plots and detailed gene counts can be found in Supplementary Figure 5. C: Differentially expressed genes that are upregulated between offset time points during slender and stumpy differentiation. The offset comparison aligns slender 15 hrs with stumpy 0 hrs, where only 22 genes are differentially expressed. Genes with an absolute log2FC > 2 and an adjusted p-value < 0.01 were classified as differentially expressed. Corresponding volcano plots and detailed gene counts can be found in Supplementary Figure 6.

Principal component analysis (PCA) was performed, with the 1^st^ principal component accounting for 32.2% and the 2^nd^ principal component for 11.1% of total variance (Figure 3A).

When comparing corresponding timepoints between slender and stumpy forms, for example, slender 8 hrs versus stumpy 8 hrs, both cell types initially exhibit a high number of differentially expressed genes. This number gradually decreases, reaching zero at 72 hrs (Figure 3B; Supplementary Figure 5). Interestingly, when comparing all time points, an additional convergence with only 22 differentially expressed genes is observed between slender 15 hrs and stumpy 0 hrs (Figure 3C; Supplementary Figure 6).

At first glance, this might suggest that slender cells transition into the stumpy form at this point before proceeding to differentiate into procyclic forms in the same way as stumpy cells. However, comparison of subsequent timepoints reveals a different pattern: at slender 24 hrs and stumpy 8 hrs, the number of differentially expressed genes increases markedly to 316. This elevated number remains relatively consistent until both cell types converge at 72 hrs, where no differentially expressed genes are detected (Figure 3B; Supplementary Figures 5 and 6).

In line with these findings, Gene Ontology (GO) enrichment analysis reveals that gene expression profiles of slender and stumpy cells throughout differentiation are associated with distinct molecular functions and biological processes. In slender cells, the most significant GO terms are related to extracellular structure and matrix organization, glycolytic processes and proteolysis (Supplementary Figure 7). In contrast, stumpy cells show enrichment for genes that are involved in RNA processing, ribosome biogenesis as well as cellular component biogenesis, consistent with them re-entering the cell cycle to become procyclic forms (Supplementary Figure 7). Transcript data of the stumpy markers PAD1 and PAD2 show different expression levels for slender and stumpy forms, before reaching similar levels by 72 hrs, when both forms had become procyclic (Supplementary Figure 8). Genes associated with the procyclic form, like the procyclin surface proteins EP1 and EP2, as well as pyruvate phosphate dikinase (PPDK), start with low expression levels for both, slender and stumpy forms. However, expression increases to procyclic levels by only 8 hrs in stumpy cells but more gradually in slender cells, reaching comparable levels only by 72 hrs (Supplementary Figure 9).

Collectively, these data indicate that differentiation in slender and stumpy trypanosomes proceeds via distinct gene expression programs. Rather than following a shared trajectory, each form activates a unique set of genes to transition into the first fly form, the procyclic form (Figure 4).

**Figure 4:**
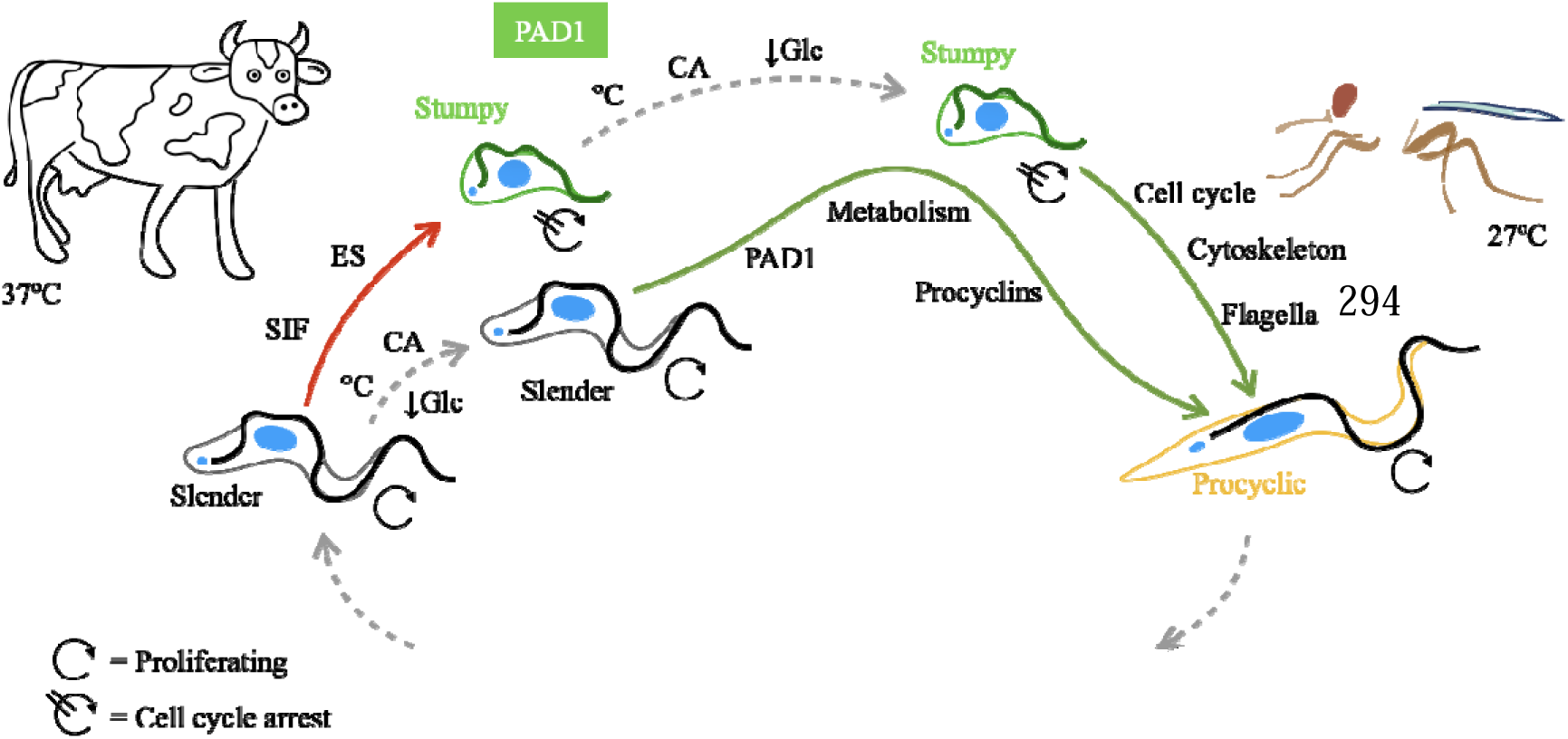
Slender and stumpy bloodstream forms must activate distinct pathways to transform into the procyclic form in the tsetse fly. Differentiation to the PAD1-positive (green), cell cycle-arrested, short stumpy form can be triggered by either SIF or ES-attenuation(Zimmermann et al., 2017). Stumpy forms have a 2-3 day window to be ingested by a tsetse fly before they perish. When a tsetse fly takes a blood meal, it can ingest both slender and stumpy forms. Once in the fly’s midgut, both forms begin their transformation into the procyclic fly form. During the initial 15 hrs, slender forms shift towards stumpy gene expression before diverging again. Stumpy forms need to reactivate the cell cycle, fully switch to proline metabolism, and elongate both their cytoskeleton and flagella (Supplementary Figure 7A). Slender forms must activate the essential PAD1 pathway, complete the switch to proline metabolism, and change to a procyclin coat (Supplementary Figure 7B). By 72 hrs into differentiation, both slender and stumpy forms have transitioned into the procyclic form. This Figure was adapted from Schuster *et al*. 2021.

## Discussion

Historically, the lack of molecular markers to distinguish slender and stumpy forms resulted in conflicting reports on their transmissibility to the tsetse fly. Early experiments indicated that higher trypanosome levels in mammalian blood correlated with more infected flies. Since stumpy forms arise through a density-dependent mechanism, increased bloodstream concentrations inevitably meant more stumpy forms in mammalian hosts, an uncontrollable factor in these early studies (Robertson, 1912; Van Hoof, 1947; Baker & Robertson, 1957; Wijers, 1958).

Moreover, the discovery of the secreted quorum sensing factor SIF in the 1990s reinforced the notion that this pathway serves to ensure the presence of stumpy forms (Reuner et al., 1997; Vassella et al., 1997). It was also shown that stumpy forms begin developing traits needed for survival in the fly midgut, such as an elaborated mitochondrion and expressing enzymes associated with the Krebs cycle (Vickerman, 1965; Brown et al., 1973; Hamm et al., 1990; Reuner et al., 1997). Previous research also suggested that the stumpy form was more adapted to life in the tsetse fly, pointing to their resistance to proteolytic stress and acidic conditions *in vitro* (Nolan et al., 2000). Later, however, it was shown that the tsetse midgut is actually an alkaline environment, making the stumpy resistance to acidic conditions mute in the case of midgut survival (Liniger et al., 2003).

Taken together, it is understandable why early studies concluded that only stumpy form trypanosomes could infect tsetse flies. This assumption, however, gave rise to the transmission paradox: under conditions of low parasitaemia during chronic infections, how can sufficient numbers of stumpy forms be maintained to ensure parasite transmission and sustain the disease cycle?

The ability to study the differences between slender and stumpy cells changed upon the discovery of the first stumpy specific molecular marker, the protein associated with differentiation 1 (PAD1), in 2009 (Dean et al., 2009). Using the PAD1 marker, it was shown that skin-resident trypanosomes reveal a locally increased proportion of stumpy forms, providing a possible solution to this transmission paradox. In mice, the proportion of stumpy cells in the skin ranged from 8 - 80%, suggesting that these parasites might significantly contribute to infection dynamics (Capewell et al., 2016).

Our recent study using artificial human skin models revealed that trypanosomes freshly deposited into the skin by tsetse flies develop into a quiescent form that cannot reinfect the vector again (Reuter et al., 2023). Although this finding does not address parasites that migrate back into the skin from circulation, it highlights the complex dynamics occurring within tissue. Ultimately, the extent to which skin-resident populations contribute to tsetse transmission remains unclear, as direct quantification currently is not feasible.

In 2021, we offered an alternative explanation for the transmission paradox, namely that slender form trypanosomes can also infect and complete the life cycle in tsetse flies. All bloodstream form trypanosomes taken up in a bloodmeal have the potential to infect a mammalian host by establishing salivary gland infection in tsetse flies (Schuster et al., 2021). While all our data supported this conclusion, some questions were raised, which we have now answered.

To mimic more natural conditions, we conducted infection experiments using slender forms without immune-suppressors, both male and female flies, and non-teneral flies.

The definition of a non-teneral and teneral fly is of great importance here.

The age of a teneral fly ranges anywhere from 8 to 72 hours post eclosion (hpe), with the important criterion that they have not taken up any blood (Otieno et al., 1983; Walshe et al., 2011). Thus, when infecting teneral flies, their first bloodmeal is the infectious one. In contrast, we infected our non-teneral flies between 144 - 168 hpe, having received two regular bloodmeals, each two days apart, before the third infectious feeding.

While it could be argued that it would be better to use even older flies, the lifespan of the tsetse must be taken into consideration to ensure survival through the 30 days required for *T. brucei* to complete development in the vector. It has been suggested that tsetse flies in the wild may live longer than their laboratory raised counterparts, though estimated and observed life spans vary widely. The mean life span ranges from 35 to 178 days for female, and only 21 to 28 days for male flies (Jackson, 1949; Maudlin et al., 1999; Vale & Torr, 2005; Haines et al., 2020). Regardless of the individual lifespan of a fly, a trypanosome infection is permanent.

We have demonstrated, in a statistically well-controlled manner, that a single slender trypanosome is capable of infecting the tsetse fly. Here, we unambiguously show that, in the absence of immunosuppressive treatment, slender forms can establish infections in tsetse flies, irrespective of the fly’s age or sex.

The study by Ngoune *et al*. (2025) does not disprove this finding (Ngoune et al., 2025). In their infection experiments, only 63% of the “stumpy” cells expressed PAD1, meaning that 37% must, by definition, have been slender forms. This heterogeneity complicates the interpretation of their results. It is also worth noting that Ngoune *et al*. used the AnTat 1.1E strain, which they maintained in culture without methylcellulose. We have observed that, during adaptation to methylcellulose-free medium, our *bona fide* pleomorphic AnTat 1.1 strain gradually loses its developmental competence. This could account for the heterogenous “stumpy” population reported by Ngoune *et al*.

Moreover, more than 40% of the adult male flies in their study did not survive the 28-day period required before dissection, further complicating the interpretation of their infection data.

In response to our original work Matthews & Larcombe postulated that “molecular characteristics” (PAD1 in particular) define a stumpy cell and that the eponymous morphological changes (e.g. stumpy formation) are no longer valid criteria (Matthews & Larcombe, 2022). Although irreversible cell cycle arrest is still an accepted hallmark of stumpy trypanosomes, we cannot exclude that in the tsetse fly, the dividing slender population arrests the cycle and then transitions to the procyclic form. This scenario is of some theoretical interest, especially in view of the ongoing discussion about shallow vs. deep cell cycle arrest, e.g. in stem cell quiescence (Urbaìn & Cheung, 2021). Therefore, we decided to conduct an RNA sequencing time course that would compare the transcriptional landscape of slender and stumpy bloodstream populations during differentiation to the procyclic form (Figure 3). Using fluorescence activated cell sorting, slender cultures were sorted to exclude any PAD1-positive cells, while stumpy cultures were sorted to include 100% PAD1 expressing cells.

At the outset, more than 300 genes were differentially expressed between the two bloodstream forms. However, after 72 hours (hrs) under differentiation conditions, no significant differences in gene expression remained; both slender and stumpy cells had transitioned into procyclic forms. Importantly, stumpy forms appear transcriptionally primed for rapid progression, while slender cells activate a distinct and temporally delayed gene expression program (Supplementary Figure 9).

The transient similarity observed at the specific timepoint, slender 15 hrs and stumpy 0 hrs, may reflect a brief convergence in gene expression, but is not indicative of a shared developmental pathway. Instead, the sustained differences in transcriptional profiles and enriched GO terms (Supplementary Figures 5, 6 and 7) argue for fundamentally different regulatory mechanisms underlying the commitment to procyclic differentiation.

These data argue against a linear progression from slender to stumpy to procyclic. Instead, they suggest that slender forms transiently activate a subset of genes also expressed in stumpy cells but then follow a distinct transcriptional trajectory before ultimately converging with the stumpy pathway at 72 hrs, when both forms adopt a procyclic identity (Figure 3). Thus, slender and stumpy forms remain transcriptionally distinct for at least the first 24 hrs of differentiation (Figure 3B). At this point, their transcriptomes do not correspond clearly to any of the three canonical forms - slender, stumpy, or procyclic. These findings demonstrate that slender forms can differentiate directly into procyclic forms without passing through a *bona fide* stumpy stage, i.e. without cell cycle arrest.

If slender forms can adopt a distinct transcriptional trajectory towards the procyclic state *in vitro*, there is no reason to assume that *T. brucei* could not use both routes - via stumpy or directly from slender - in the tsetse fly as well.

Additionally, there is one more observation that needs to be taken into account. We have shown that even small numbers of monomorphic trypanosomes strain 427 - incapable of differentiating into stumpy forms - can establish stable midgut infections in tsetse flies yet fail to progress to salivary gland colonization (Schuster et al., 2021). These monomorphic cells do not respond to Stumpy Induction Factor (Reuner et al., 1997; Vassella et al., 1997) and fail to upregulate PAD1 (Dean et al., 2009), remaining locked in the proliferative slender bloodstream form. Thus, slender trypanosomes are indeed capable of infecting the tsetse midgut, even if they are monomorphic. The PAD pathway is not strictly required for colonization of the midgut. However, activation of the PAD pathway is essential for the generation of procyclic trypanosomes that are competent to complete development and colonize the salivary glands.

Our results clearly support the conclusion that slender bloodstream forms can successfully establish infection in the tsetse fly (Schuster et al., 2021), and extend the findings of Larcombe et al. (2023), who proposed that transmission during high parasitaemia is predominantly mediated by stumpy forms and pre-committed slender intermediates.

Notably, according to the criteria summarized in Larcombe et al.’s developmental commitment classification (Larcombe et al., 2023; Figure 4A), the populations used in our study qualify as “true slender” cells, defined as replicative forms that neither express PAD1 nor display stumpy morphology. In our infection experiments, these sorted slender cells continued proliferating and did not arrest the cell cyle at the time of fly infection (Supplementary Figure 3C), confirming their identity as “true slender” forms under Larcombe et al.’s definition.

This underscores that our transmission data were generated using precisely the cell type previously proposed to be both rare and transmission-incompetent.

Consequently, our findings resolve the apparent transmission paradox: while stumpy and intermediate forms may dominate at high parasitaemia (Larcombe et al., 2023), slender cells prevail during low-parasitaemia phases and are themselves sufficient to maintain the *T. brucei* life cycle through transmission. Thus, replicative slender form should be recognized as a *bona fide* transmission stage, not merely a proliferative bloodstream form. Therefore, in retrospect, our results are perhaps less unexpected than initially assumed.

## Material and Methods

### Cell line

The pleomorphic *Trypanosoma brucei brucei* strain EATRO 1125 (serodome AnTat 1.1) (Le Ray et al., 1977) with an NLS-GFP PAD1 3’UTR molecular marker and an additional tdTomato fluorescence sequence (Reuter et al., 2023) was used for infection experiments and cultured as previously described (Schuster et al., 2021). For RNA sequencing experiments the same cell line was used with an additional EP1:YFP fusion protein (Schuster et al., 2021).

### Fly infection

Tsetse flies of the species *Glossina morsitans morsitans* were kept as previously described (Schuster et al., 2021).

Teneral flies received their first, and infectious, bloodmeal 24-72 hours post-eclosion (hpe). They were infected with four trypanosomes per bloodmeal, either untreated or supplemented with 60 mM *N*-acetyl-glucosamine (NAG).

Non-teneral flies (144-168 hpe) were given two non-infectious bloodmeals, each two days apart, prior to the infectious bloodmeal containing 1×10^6^ cells/ml of either slender or stumpy parasites. Prior to infection, fluorescence activated cell sorting was performed to ensure 100% slender or stumpy population.

### Immunofluorescence

Cells were prepared, fixed and stained as previously described (Schuster et al., 2021).

### Fluorescence activated cell sorting

Cell sorting was performed using the FACS Aria III (BD biosciences, Franklin Lakes, USA). Cells were harvested, resuspended in pre-warmed TDB at 1×10^7^ cells/ml, transferred to FACS tubes through a 35 µm cell strainer cap. Cells were first gated based on the tdTomato signal to exclude debris and residual medium. Subsequently, GFP fluorescence from the NLS-GFP PAD1 3’UTR reporter was analysed. Depending on the desired population, gates were set to isolate either PAD1-positive (stumpy) or PAD1-negative (slender) cells with high purity. For each experiment, 1×10^6^ cells were sorted using a 100 µm nozzle at the lowest sorting speed to minimize mechanical stress. Sorted cells were collected into 15 ml tubes containing pre-warmed FCS, resulting in a final FCS concentration of 15% (v/v). After sorting, cell motility, concentration, and PAD1 signal were checked by microscopy.

### Bulk RNA sequencing

Both slender and stumpy trypanosomes were differentiated into procyclic forms as previously described. The 0 hour (hr) timepoint was collected immediately before differentiation was induced, and additional samples were taken at 8, 15, 24, 72 hrs following addition of *cis*-aconitate.

Cells were harvested from biological triplicates, resuspended in 1 ml pre-warmed PBS containing 10 % FCS, and transported to the Helmholtz Institute for RNA Infection biology (HIRI) Würzburg in a 37°C incubation chamber.

Upon arrival, cells were washed twice with pre-warmed PBS to remove FCS in 1 ml of pre-warmed PBS 15 min prior to sorting, Calcein-AM violet was added at a final concentration of 1 µM to label viable cells.

Live cells were sorted using a FACS Aria III cytometer (BD Biosciences). 1000 cells per replicate were deposited into single wells of a 48 well-plate containing 2.6 µl of 1x lysis buffer (Takara) and 0.01 µl of RNase inhibitor (40 U/µl; Takara). Sorting was performed in triplicates for each timepoint, and plates were immediately placed on ice and stored at -80°C. Library preparation and sequencing were performed as previously described (Müller et al., 2018). Very briefly, lysates were supplemented with 0.2 µl ERCC Spike-in Control Mix 1 (Thermo Fisher Scientific) at a 1:20,000,000 dilution. Libraries were prepared using the SMART-Seq v.4 Ultra Low Input RNA Kit (Takara), utilizing a quarter of the recommended reagent volumes. PCR amplification was performed for 27 cycles, and cDNA was purified with Agencourt AMPure XP beads (Beckman Coulter) and 15 µl elution buffer (Takara). Library quantification was carried out using a Qubit 3 Fluorometer and dsDNA Hs Assay kit (Life Technologies), while quality assessment was performed by using a 2100 Bioanalyzer with High Sensitivity DNA kit (Agilent). 0.5 ng of cDNA was used as input for the Nextera XT (Illumina) tagmentation-based library preparation protocol. The reaction was performed at one quarter of the recommended volumes, with a 10-minute tagmentation step at 55 °C, and a 1-minute extension step during PCR. Libraries were pooled and sequenced in paired-end mode with 2x 75 cycles using Illumina’s NextSeq 500.

### Analysis of bulk RNA-sequencing data

Bulk RNA-sequencing analysis was performed according to scripts and online resources published by Berry *et al*., 2021 using RStudio (version 4.02). Read demultiplexing and quality control were performed with FASTQC (version 0.12.1, (Andrews, 2012)). Adaptors were removed and reads were trimmed using Fastp (version 0.23.4, (Chen, 2023)).

The *T. b. brucei* TREU 972 reference genome was obtained from the Tritryp database (TritrypDB). rRNA was removed (Aslett et al., 2010), and, to remove redundancy, only one representative from each gene group was retained while others were masked. Gene groups were defined based on sequence similarity using CD-Hit software (Li & Godzik, 2006).

Kallisto was used for read alignment, and genes with ≥ 3 counts per million (cpm) in at least two samples were retained for downstream analysis. Data normalization was performed using the EdgeR package in RStudio (Robinson et al., 2009; Team R.C., 2021).

SL152 (slender 15 hrs, 2^nd^ replicate) and SL241 (24 hrs, 1^st^ replicate) were considered outliers based on hierarchical clustering distance using the Minkowski metric (Lee & Willcox, 2014). Differentially expressed genes (DEGs) were identified using DESeq2, EdgeR, and Limma packages in RStudio (Love et al., 2014; Ritchie et al., 2015; Robinson et al., 2009; Team R.C., 2020), applying significance criteria of log2 fold change > 1 and p value < 0.01.

### Statistics

Two-tailed Fisher’s exact tests were performed for all fly infection data using GraphPad prism version 9.0.0 for Windows (Graphpad software, San Diego, California USA, http://www.graphpad.com).

## Acknowledgements

The authors would like to thank the fly team of the Zoology I Department of the University of Würzburg for their expert care and maintenance of the tsetse flies.

