## Supplemental Figure for "Research advance: Unexpected plasticity in the life cycle of *Trypanosoma brucei*"

| No. of flies infected | Fly sex | No. of slender cells/ml | Concentration of NAG | Days until dissection | No. of flies dissected | MG | PV | SG |
| --- | --- | --- | --- | --- | --- | --- | --- | --- |
| 37 | male | 200 | 0 mM | 35 | 16 | 0 | 0 | 0 |
| 19 | female | 200 | 0 mM | 35 | 17 | 1 | 1 | 0 |
| 24 | male | 200 | 0 mM | 35 | 24 | 5 | 4 | 1 |
| 24 | female | 200 | 0 mM | 35 | 22 | 2 | 2 | 0 |
| 24 | male | 200 | 0 mM | 35 | 21 | 1 | 1 | 0 |
| 9 | female | 200 | 0 mM | 35 | 9 | 1 | 1 | 0 |

| No. of flies infected | Fly sex | No. of slender cells/ml | Concentration of NAG | Days until dissection | No. of flies dissected | MG | PV | SG |
| --- | --- | --- | --- | --- | --- | --- | --- | --- |
| 37 | male | 200 | 60 mM | 35 | 35 | 3 | 3 | 2 |
| 19 | female | 200 | 60 mM | 35 | 18 | 1 | 1 | 0 |
| 26 | male | 200 | 60 mM | 35 | 24 | 1 | 1 | 1 |
| 27 | female | 200 | 60 mM | 35 | 27 | 1 | 1 | 0 |
| 26 | male | 200 | 60 mM | 35 | 26 | 7 | 4 | 2 |
| 9 | female | 200 | 60 mM | 35 | 8 | 3 | 3 | 0 |

Supplementary Figure 1: Absolute numbers of tsetse fly infections using slender bloodstream forms of *T. brucei*, with or without the addition of *N*-Acetyl-Glucosamine (NAG). Both, male and female flies were infected with blood containing 200 slender cells per ml of blood, either untreated or supplemented with the immune-suppressing chemical, *N*-Acetyl-Glucosamine (NAG, 60mM). Tsetse flies have an estimated drinking volume of 20 µl (Gibson & Bailey, 2003), which results in an uptake of 4 parasites per bloodmeal. All flies were dissected 35 days post infection, and their midgut (MG), proventriculus (PV), and salivary glands (SG) examined for parasite presence. Prior to infection, slender cells of the tdTomato NLS-GFP:PAD1 3´UTR line were verified to lack *pad1* expression, confirming pure slender identity (< 0.05% PAD1 positive).

| No. of flies infected | Fly sex | No. of slender cells/ml | Concentration of NAG | Days until dissection | infectious feed | | No. of flies dissected | MG | PV | SG |
| --- | --- | --- | --- | --- | --- | --- | --- | --- | --- | --- |
| 20 | male | 1x10^6^ | 0 mM | 30 | 3^rd^ | 17 | | 0 | 0 | 0 |
| 20 | male | 1x10^6^ | 0 mM | 30 | 3^rd^ | 19 | | 2 | 2 | 2 |
| 21 | male | 1x10^6^ | 0 mM | 35 | 3^rd^ | 21 | | 5 | 4 | 2 |
| 20 | male | 1x10^6^ | 0 mM | 35 | 3^rd^ | 19 | | 0 | 0 | 0 |
| 20 | female | 1x10^6^ | 0 mM | 30 | 3^rd^ | 20 | | 1 | 0 | 0 |
| 21 | female | 1x10^6^ | 0 mM | 30 | 3^rd^ | 21 | | 2 | 2 | 2 |
| 20 | female | 1x10^6^ | 0 mM | 35 | 3^rd^ | 20 | | 0 | 0 | 0 |
| 20 | female | 1x10^6^ | 0 mM | 35 | 3^rd^ | 20 | | 0 | 0 | 0 |

| No. of flies infected | Fly sex | No. of stumpy cells/ml | Concentration of NAG | Days until dissection | infectious feed | No. of flies dissected | MG | PV | SG |
| --- | --- | --- | --- | --- | --- | --- | --- | --- | --- |
| 20 | male | 1x10^6^ | 0 mM | 38 | 3^rd^ | 20 | 2 | 2 | 1 |
| 21 | male | 1x10^6^ | 0 mM | 36 | 3^rd^ | 16 | 1 | 1 | 1 |
| 23 | male | 1x10^6^ | 0 mM | 35 | 3^rd^ | 19 | 2 | 2 | 0 |
| 14 | male | 1x10^6^ | 0 mM | 35 | 3^rd^ | 11 | 0 | 0 | 0 |
| 20 | female | 1x10^6^ | 0 mM | 38 | 3^rd^ | 19 | 1 | 0 | 0 |
| 20 | female | 1x10^6^ | 0 mM | 36 | 3^rd^ | 20 | 1 | 0 | 0 |
| 23 | female | 1x10^6^ | 0 mM | 35 | 3^rd^ | 19 | 0 | 0 | 0 |
| 15 | female | 1x10^6^ | 0 mM | 35 | 3^rd^ | 13 | 2 | 1 | 0 |

Supplementary Figure 2: Absolute numbers of infections in non-teneral tsetse flies with either slender or stumpy *T. brucei* cells. Male and female flies were infected with untreated blood containing 1x10^6^ cells/ml. The tdTomato NLS-GFP:PAD1 3´UTR cell line enabled FACS-based separation of stumpy (PAD1 positive*,* GFP in nucleus) and slender (PAD1 negative, no nuclear fluorescence) forms prior to infection. All non-teneral flies were 144-168 hours post eclosion (hpe) and had already received two non-infectious bloodmeals prior to the infectious feed. Flies were dissected 35 days post infection, and their midgut (MG), proventriculus (PV), and salivary glands (SG) examined for parasite presence.

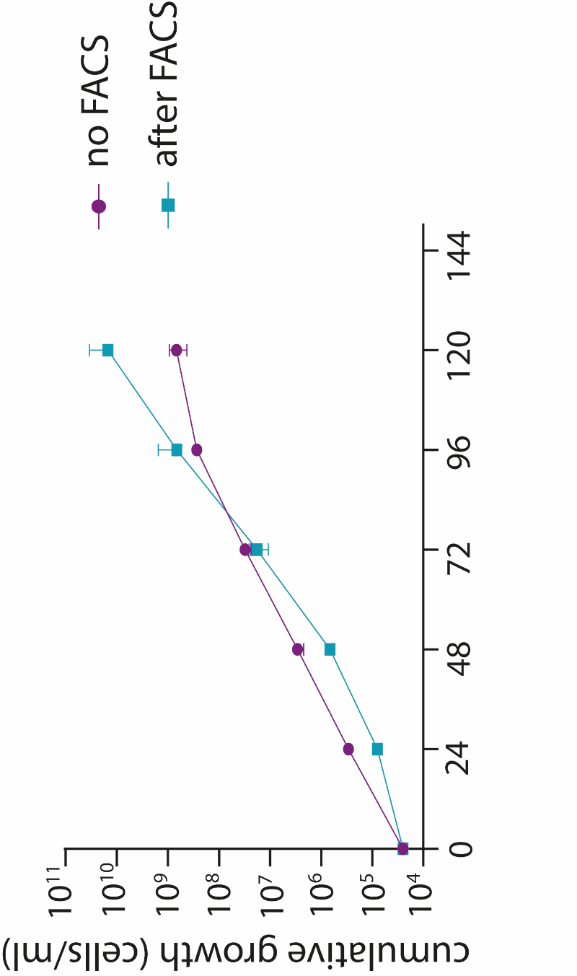

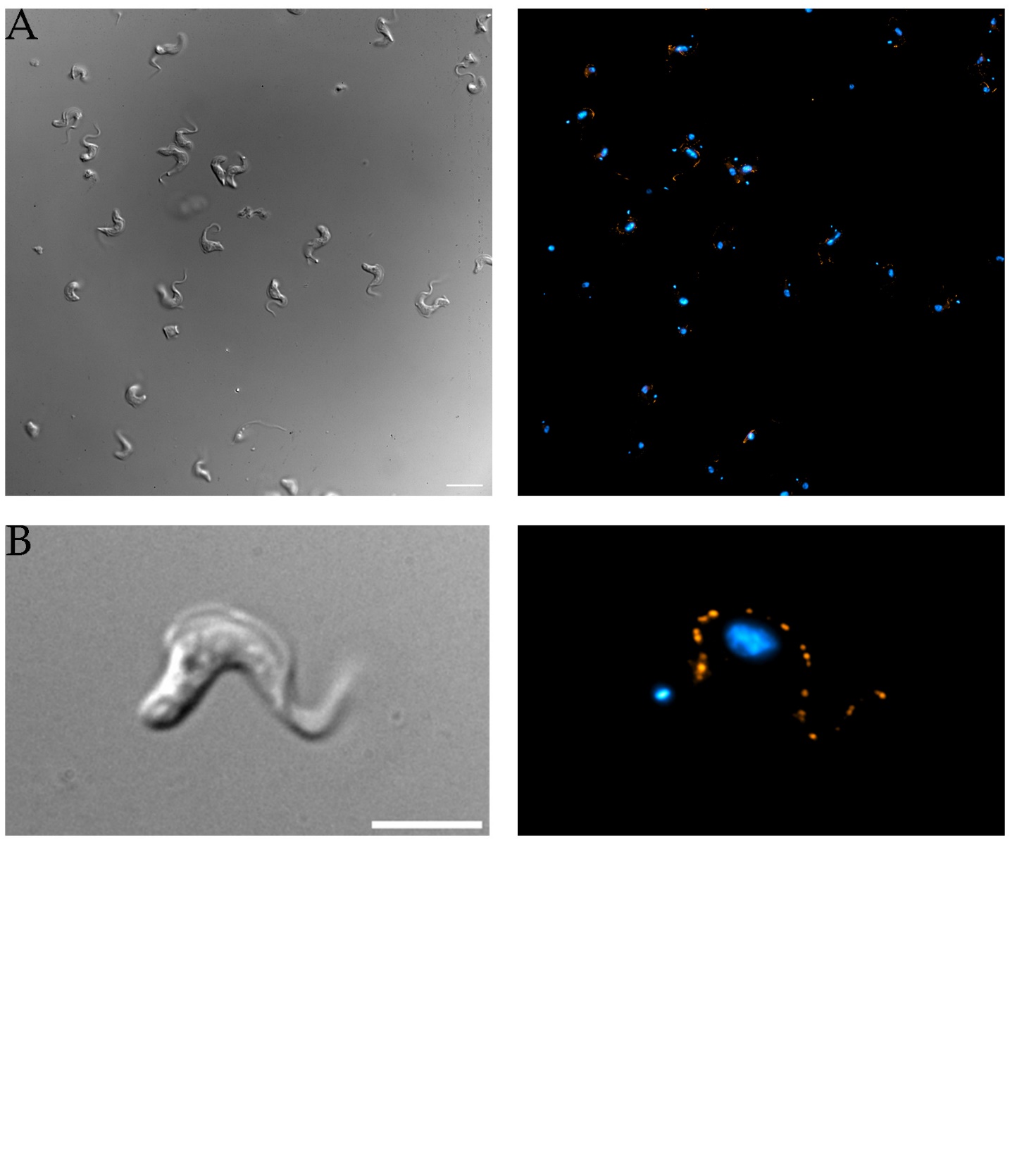

C

Supplementary Figure 3: Slender *T. brucei* cells of the tdTomato NLS-GFP:PAD1 3´UTR line do not express PAD1 or exhibit stress responses following fluorescence activated cell sorting (FACS). FACS was used to ensure pure slender populations (*pad1* negative) prior to infection. A: Immunofluorescence (IF) images of slender cells immediately after FACS, confirming sorting success. Parasites were fixed in 4% Paraformaldehyde (PFA), stained with DAPI (blue), and labelled with an anti-PAD1 antibody (orange); scalebar: 20 µm. B: High-resolution IF image of a single slender trypanosome post sorting, showing clear absence of nuclear *pad1* signal; scalebar: 10 µm. C: Growth curves comparing slender cells post sorting (green) and untreated slender cells (purple), indicating no growth impairment due to sorting.

Supplementary Figure 4: Fluorescence activated cell sorting (FACS) was used to isolate stumpy cells of the tdTomato NLS-GFP:PAD1 3´UTR line, ensuring a pure stumpy population prior to infection. A: Stumpy cells from a SIF-induced stumpy culture (grown to 5x10^5^ cells/ml and kept for 48 hours), after FACS to confirm sorting success. Cells were fixed in 4% PFA immediately after sorting, stained with DAPI (blue), and labelled with an anti-PAD1 antibody (orange); scalebar: 20 µm.
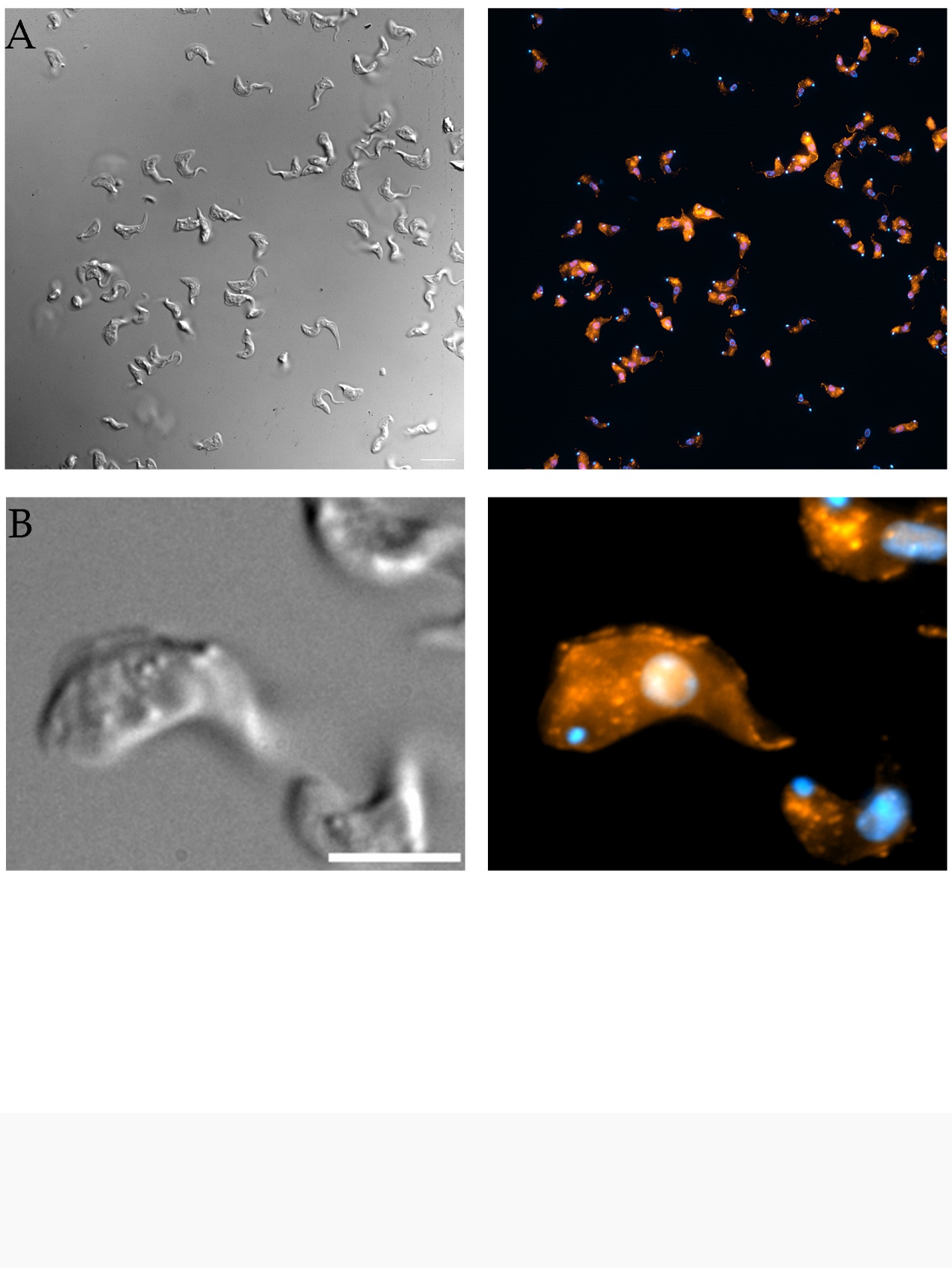
 B: High-resolution IF image of a single stumpy trypanosome displaying characteristic stumpy morphology and strong *pad1* signal (orange); scalebar: 10 µm.

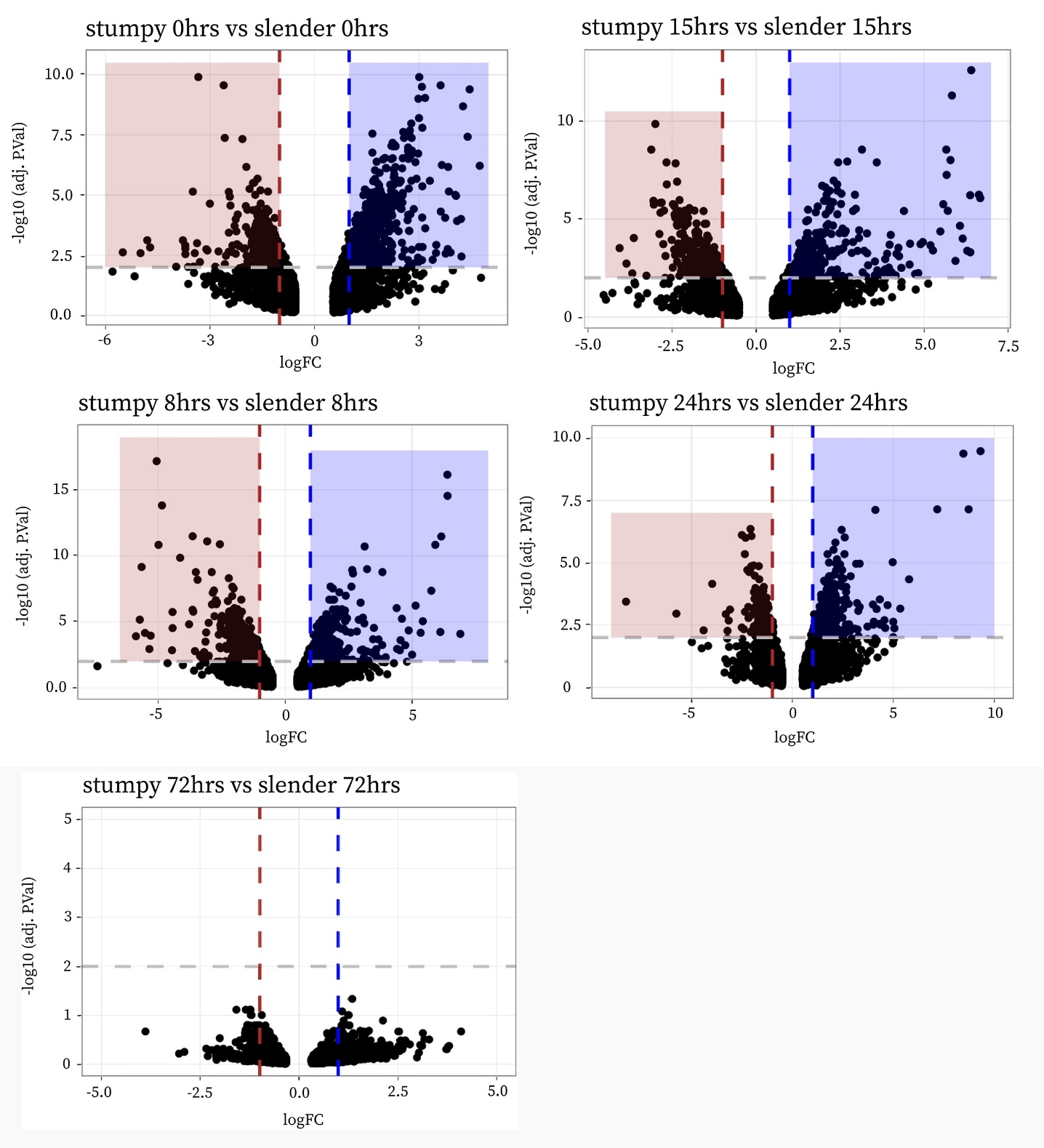

|  | slender | stumpy |
| --- | --- | --- |
| 0hrs | 533 | 314 |
| 8hrs | 310 | 362 |
| 15hrs | 325 | 292 |
| 24hrs | 272 | 175 |
| 72hrs | 0 | 0 |

Supplementary Figure 5: Volcano plots showing differential gene expression between stumpy (red) and slender (blue) *T. brucei* forms during *in vitro* differentiation to the procyclic form at 0, 8, 15, 24 and 72 hrs after induction. Differentiation was initiated by adding *cis*-aconitate, lowering the temperature to 27°C, and depleting glucose in the medium. Each black dot represents one gene. Genes within the coloured boxes show 2x up-regulation in slender (blue dotted line) or 2x up-regulation in stumpy (red dotted line) cells, with a p- value $\leq$ 0.01 (grey dotted line). Red and blue boxes highlight significantly upregulated genes in stumpy or slender, respectively, with the exact gene counts listed in the accompanying table. logFC= log2 Fold Change; hrs = hours after the addition of *cis*-aconitate.

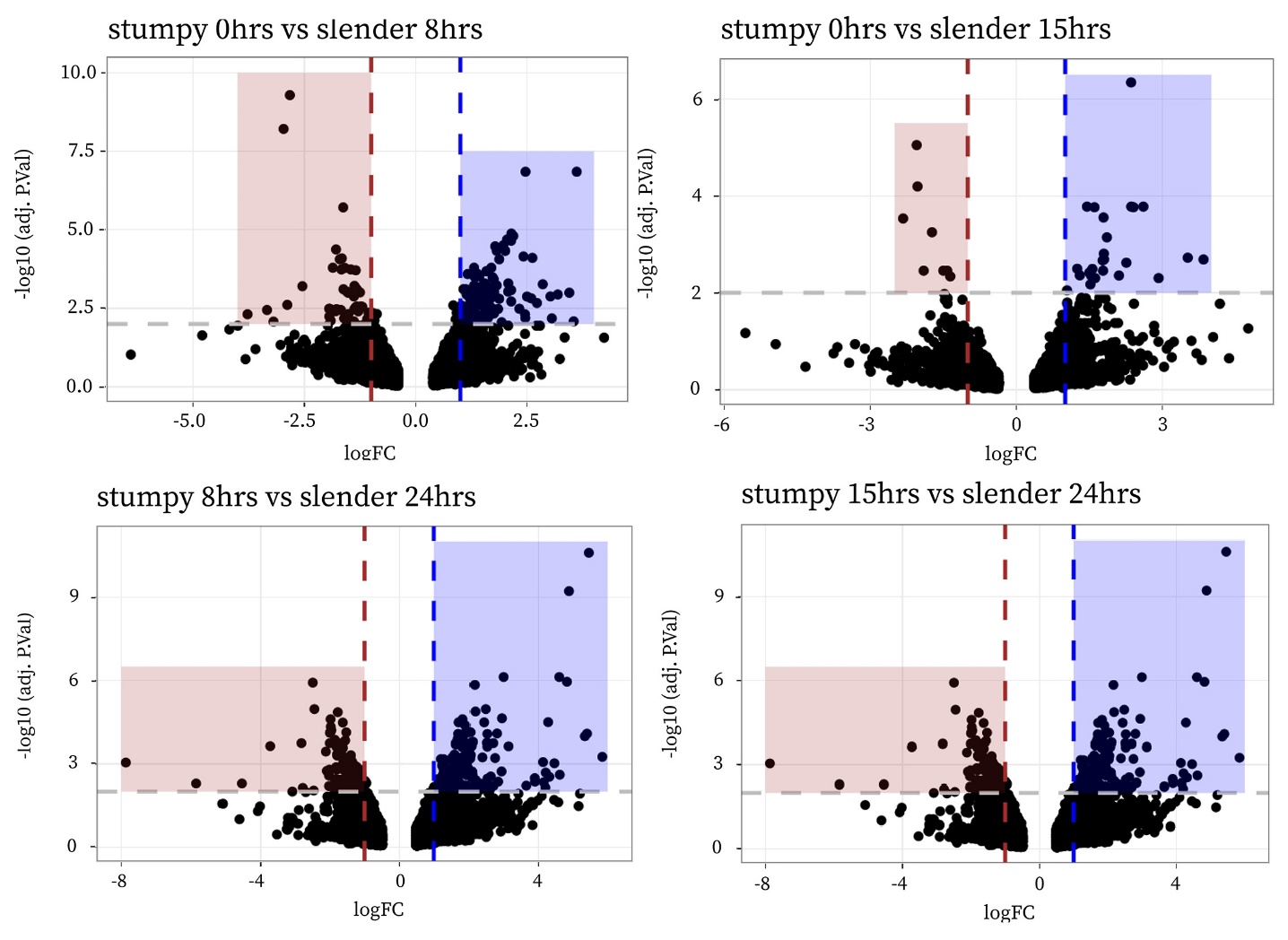

|  | slender | stumpy |
| --- | --- | --- |
| stumpy 0hrs vs slender 8hrs | 135 | 50 |
| stumpy 0hrs vs slender 15hrs | 15 | 7 |
| stumpy 8hrs vs slender 24hrs | 178 | 138 |
| stumpy 15hrs vs slender 24hrs | 178 | 138 |

Supplementary Figure 6: Volcano plots showing differential gene expression of stumpy (red) and slender (blue) *T. brucei* forms following *in vitro* differentiation to the procyclic form. Differentiation was initiated by adding *cis*-aconitate, lowering the temperature to 27°C, and depleting glucose. An offset comparison – based on proximity in the PCA plot (Figure 3A) - aligns slender cells at 15 hours (hrs) with stumpy cells at 0 hrs. Each black dot represents one gene. Genes within the coloured boxes show 2x up-regulation for slender (blue dotted line) or 2x up-regulation for stumpy (red dotted line) cells, with a p-value$\leq$ 0.01 (grey dotted line). Red and blue boxes highlight significantly upregulated genes in stumpy or slender, respectively, with the exact numbers of differentially expressed genes listed in the accompanying table. Notably, slender cells at 15 hrs exhibit a similar gene expression profile to that of stumpy trypanosomes after 0 hrs, before diverging again at later time points. logFC= log2 Fold Change; hrs = hours after the addition of *cis*-aconitate.

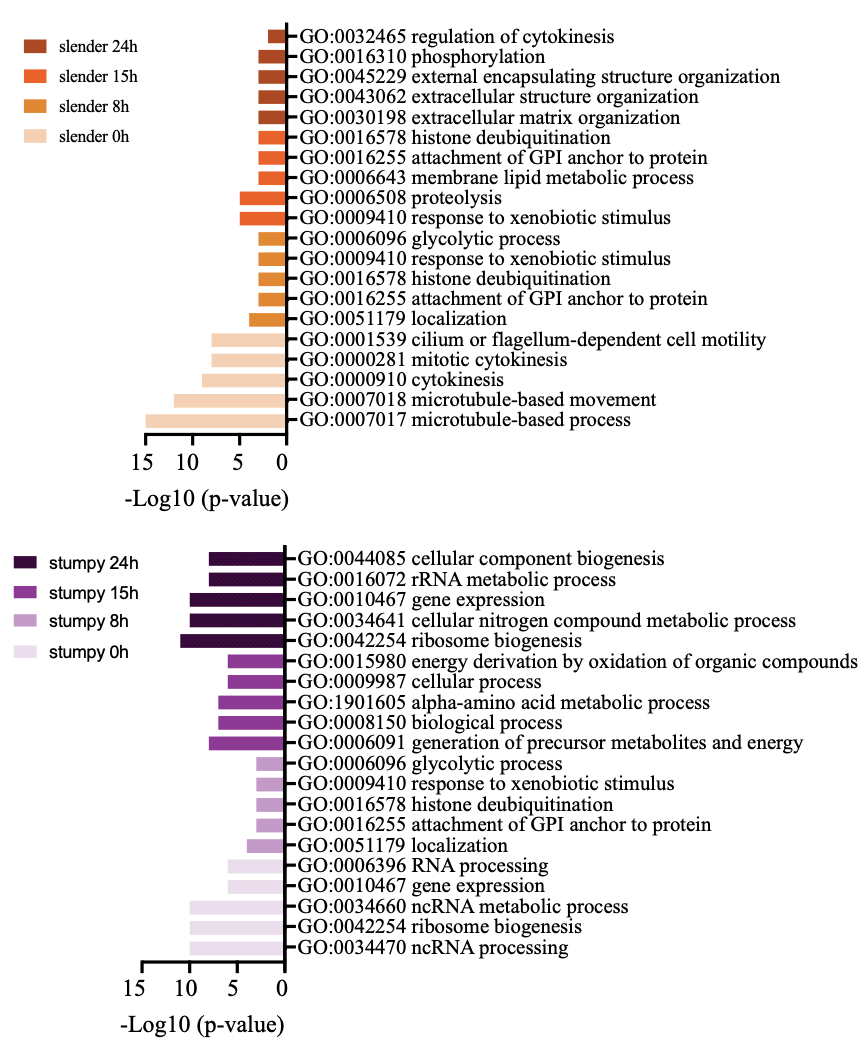

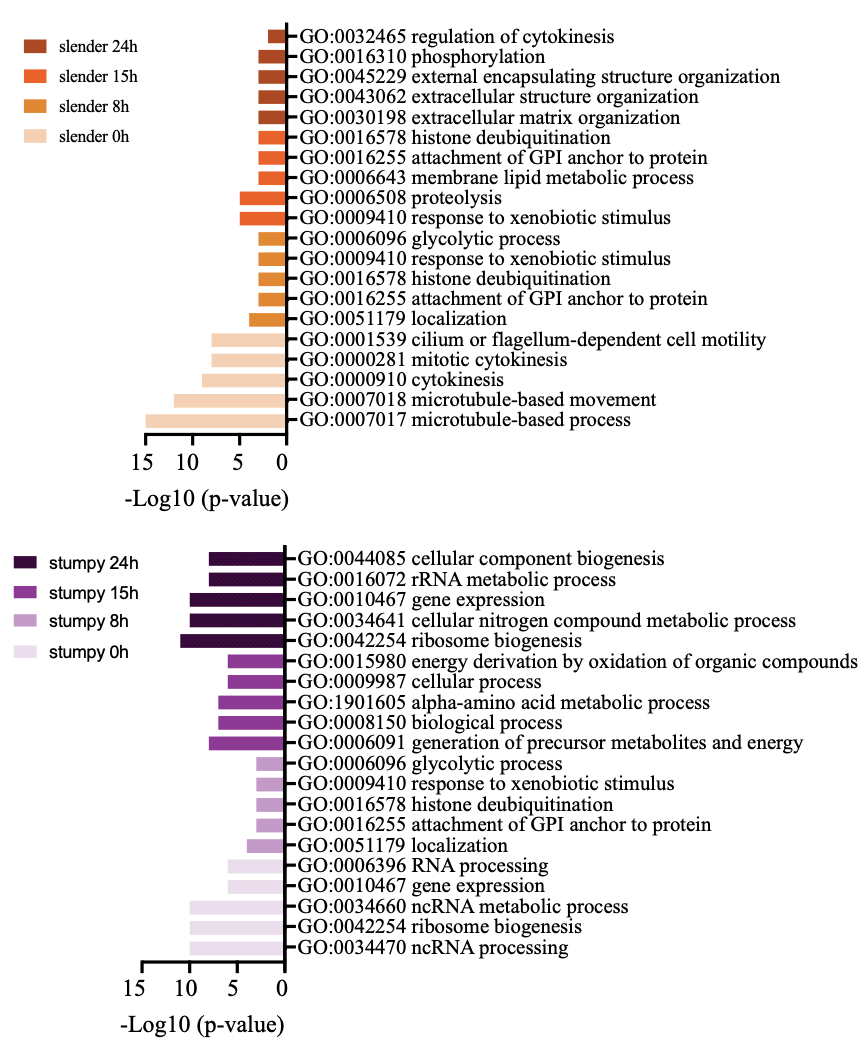

B

A

Supplementary Figure 7: During differentiation into the procyclic form, slender and stumpy parasites exhibit gene expression profiles associated with distinct biological processes and molecular functions. Gene Ontology (GO) enrichment analysis between corresponding time points identified genes with a log2 fold change of at least 1 (indicating 2x expression) and a p-value of ≤ 0.01 for either slender (orange) or stumpy (purple) forms at 0, 8, 15, 24, and 72 hours (Supplementary Figure 5). GO annotations were sourced from the TriTryp.org database (TriTryp.org) and refined using Revigo. The most significantly enriched GO terms for each time point are shown.

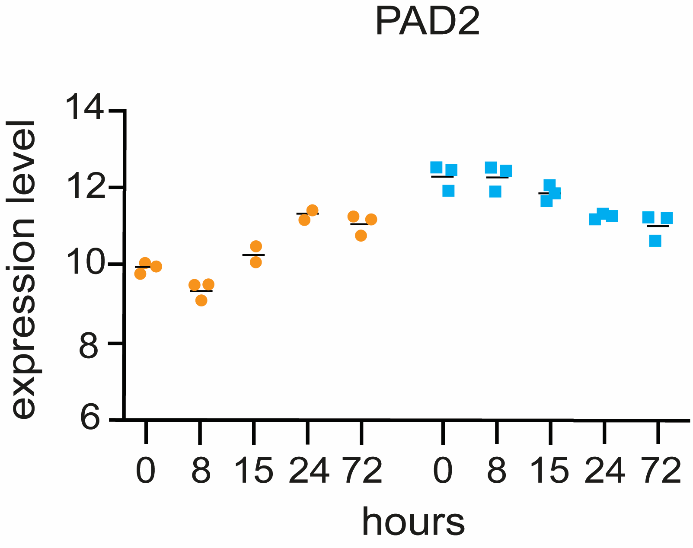

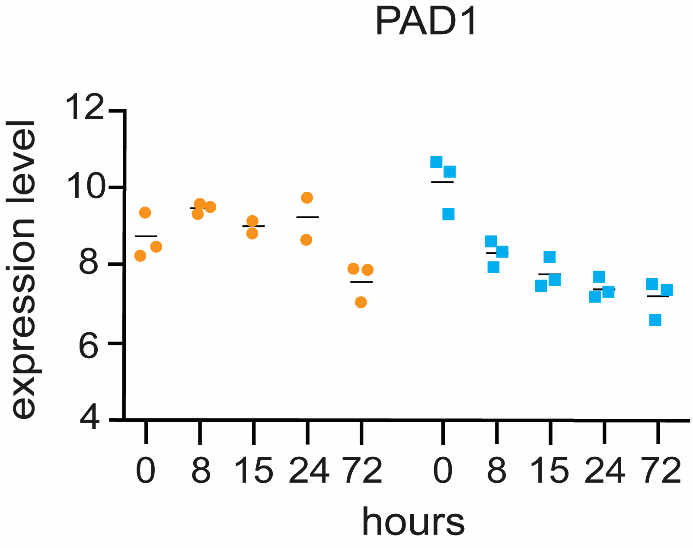

A

B

Supplementary Figure 8: Dot plots showing expression levels of genes associated with the stumpy form – PAD1 (A) and PAD2 (B) – in slender (orange) and stumpy (blue) forms during *in vitro* differentiation. Differentiation was induced by the addition of *cis*-aconitate, reduction of temperature to 27°C, and glucose depletion. RNA-sequencing was performed in triplicates; each coloured dot represents one replicate (1000 cells), and the black line indicates the mean expression value.

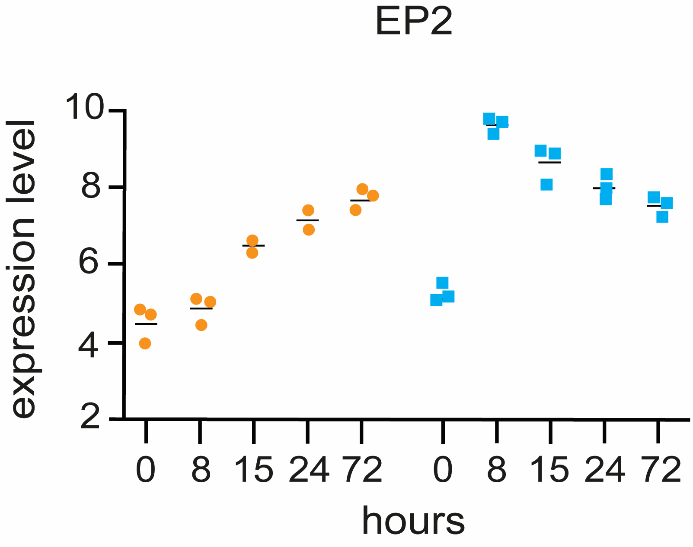

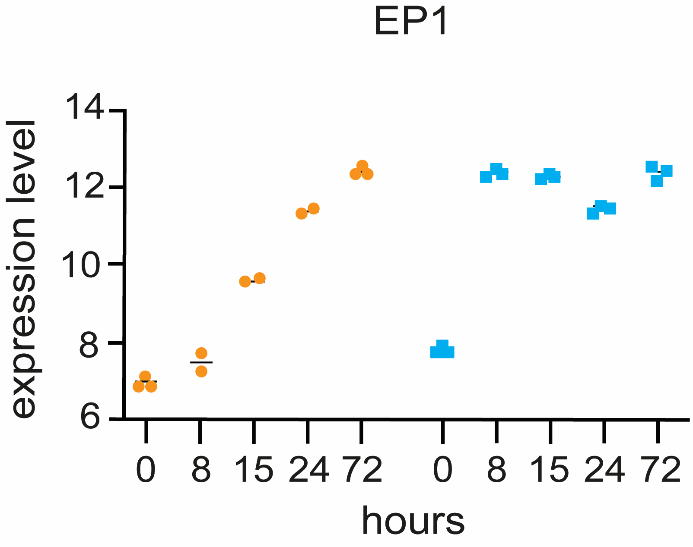

B

C

A

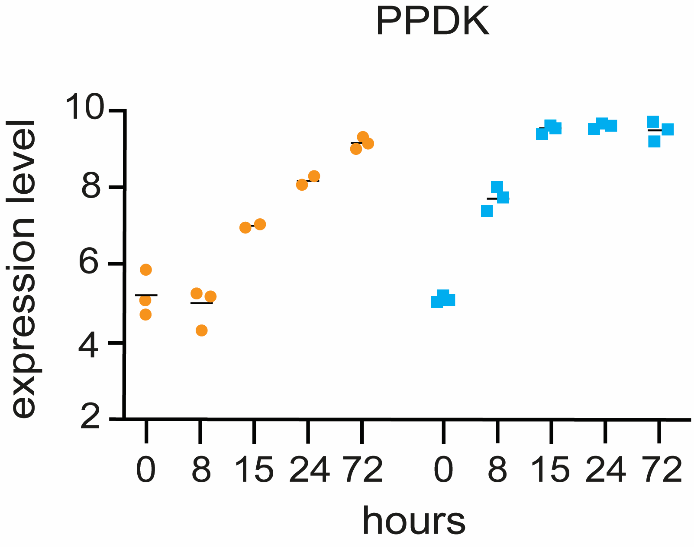
Supplementary Figure 9: Dot plots showing expression levels of genes associated with the procyclic form – EP1 (A), EP2 (B) and pyruvate phosphate dikinase (PPDK) (C) – in slender (orange) and stumpy (blue) forms during *in vitro* differentiation. Differentiation was induced by the addition of *cis*-aconitate, reduction of temperature to 27°C, and glucose depletion. Stumpy forms reach procyclic-like expression levels by approximately 8 hours, whereas slender forms display a more gradual increase. RNA-sequencing was performed in triplicates; each coloured dot represents one replicate (1000 cells), and the black line indicates the mean expression value.
